# Tuning shape and internal structure of protein droplets via biopolymer filaments

**DOI:** 10.1101/2019.12.13.876003

**Authors:** Danielle R. Scheff, Kimberly L. Weirich, Kinjal Dasbiswas, Avinash Patel, Suriyanarayanan Vaikuntanathan, Margaret L. Gardel

## Abstract

Macromolecules can phase separate to form liquid condensates, which are emerging as critical compartments in fields as diverse as intracellular organization and soft materials design. A myriad of macromolecules, including the protein FUS, form condensates which behave as isotropic liquids. Here, we investigate the influence of filament dopants on the material properties of protein liquids. We find that the short, biopolymer filaments of actin spontaneously partition into FUS droplets to form composite liquid droplets. As the concentration of the filament dopants increases, the coalescence time decreases, indicating that the dopants control the relative surface tension to viscosity. The droplet shape is tunable ranging from spherical to tactoid as the filament length or concentration is increased. We find that the tactoids are well described by a model of a bipolar liquid crystal droplet, where nematic order from the anisotropic actin filaments competes with isotropic interfacial energy from the FUS, controlling droplet shape in a size-dependent manner. Our results demonstrate a versatile approach to construct tunable, anisotropic macromolecular liquids.

## Introduction

Liquid condensates, dense macromolecular droplets that phase separate out of a dilute suspension, are ubiquitous in soft and biological materials ranging from coacervates [1] to membraneless organelles [2]. The formation and material properties of condensates can be tuned through modifying macromolecular composition [3–5] or environmental conditions [6–8]. Intriguingly, the fluid condensates typically adopt a characteristic spherical shape and coalescence, indicative of an isotropic liquid with a dominant interfacial tension. However, macromolecules are inherently structured, often with significant rigidities and size, which may impart anisotropy to the liquid.

It is well appreciated that highly anisotropic rod-like objects can form structured liquid phases [9,10]. Below a critical volume fraction, a suspension of rods is isotropic. However, above a critical packing the rods locally align due to entropic effects and adopt orientational order, forming a phase known as a nematic liquid crystal [10]. The local alignment imparts an elasticity to the fluid [11], which can depend on the packing and properties of the rods [12,13]. Liquid crystal droplets are observed to nucleate out of a dense isotropic suspension at the isotropic-nematic phase transition [14–16]. The competing effects of elasticity and surface tension in the droplets results in an elongated, spindle shape called a tactoid [17]. Due to different scaling of bulk and interfacial properties, the droplet shape is size-dependent [14,15] and is predicted to transition from spherical to tactoid [17,18]. Recently, nematic droplets that condense out of a dilute suspension with tunable shape were achieved by inducing attraction of biopolymer filaments of actin with transient cross-links [19]. The extent to which rod-like molecules influence anisotropic properties of condensates remains to be explored.

Here we create composite liquid droplets composed of FUS, a protein that forms condensates [20], doped with the biopolymer F-actin to investigate the impact of anisotropic dopants on droplet shape and properties. We find that actin incorporates throughout FUS droplets, leading to a composite liquid phase. By varying actin filament concentration and length, the degree of the anisotropic effects on the liquid vary and result in tunable droplet shape. We find that the droplet shape is well described by a continuum model of nematic droplets, where the nematic elastic energy arises from the actin filaments. Our results indicate that rigid dopants can impart liquid crystallinity to otherwise isotropic droplets. Such composite droplets provide a new means to control material properties and shape of liquid condensates, with implications for designing both biological assemblies and soft materials.

## Methods

### Protein purification

Monomeric actin (G-actin) is purified from rabbit skeletal muscle acetone powder (Pel Freeze Biologicals, Product code: 41008-3) using a procedure adapted from [21] and stored in 2 mM Tris, 0.2 mM ATP, 0.5 mM DTT, 0.1 mM CaCl_2_, 1 mM NaN_3_, pH 8. Actin is labelled using tetramethylrhodamine-6-maleimide (TMR) dye (Life Technologies). HisTag mouse capping protein is purified from bacteria using a procedure adapted from [22] and stored in a buffer composed of 10 mM Tris, 40 mM KCl, 0.5 mM DTT, 0.01 wt% NaN_3_, 50 vol% glycerol, pH 7.5. FUS-GFP is expressed in and purified from insect cells as described in [20] and stored in a buffer composed of 2 mM Tris, 500 mM KCl, 1 mM DTT, pH 7.4. All proteins are flash frozen in liquid nitrogen and stored at −80°C. Actin and capping protein are used within 3 days of thawing, while FUS is used within 4 hours. After thawing, proteins are stored at 0-4°C until use.

### Experimental assay

The experimental chamber is composed of a glass cylinder (3166-10; Corning Life Sciences) epoxied to a glass coverslip (Fisherbrand, #1.5). The coverslip surface was passivated against protein adhesion through an oil-surfactant layer. To form the layer, the surfactant, PFPE-PEG-PFPE (008; RAN Biotechnologies) is first dissolved in Novec-7500 Engineered Fluid (3M) at 2 wt%. The oil-surfactant solution is sonicated for 30 min in a bath sonicator, filtered through a 0.2 μm pore sized membrane (6784-1302; GE Healthcare), then flushed with nitrogen gas and stored at 4C until use. Coverslips are first cleaned by sonicating in ethanol, then immersed in 2 vol% triethoxy(octyl)silane (440213; Sigma-Aldrich) in isopropanol for 10 min, then submerged 24 times in water. Immediately prior to adding the sample, 4 μL of oil-surfactant solution is added to the sample chamber to create a thin layer of oil-surfactant at the coverslip. After coating the coverslip, excess solution is removed.

To polymerize actin filaments, 5 μM actin monomer (0.5 μM labelled with TMR) is added to a buffer composed of 2 mM Tris, 2 mM MgCl_2_, 25 mM KCl, 0.5 mM ATP, 0.3 wt% methylcellulose, pH 7.4. To regulate actin filament length, capping protein is added to the final concentration of 50-150 nM. Actin polymerizes into filaments for at least 30 min, before adding 0.25 μM phalloidin to prevent dilution-induced depolymerization. Actin filaments are then mixed with 4.4 μM FUS resulting in a final buffer composed of 2 mM Tris, 2 mM MgCl_2_, 45 mM KCl, 0.5 mM ATP, 0.04 mM DTT, 0.3 wt % methylcellulose, 0.1 μM phalloidin, pH 7.4. This solution is immediately added to the sample chamber. Droplets are allowed to grow for 60 min before shape is measured.

### Filament Length

We modulate the length of actin filaments through the use of capping protein, which binds to growing filaments and prevents further polymerization, leading to an exponential distribution of lengths. In the limit of strong binding, we approximate the average number of monomers in a filament to equal the ratio of actin monomers to capping protein. We convert monomers to length using the known value of ∼1 monomer per 2.7 nm in a filament. Polymerizing the actin before mixing with FUS ensures that interactions between FUS and actin do not influence actin filament polymerization and filament length. To further ensure consistent length, samples are compared only with other samples prepared on the same day when measuring the effect of actin and capping protein concentration. For varying actin concentration, all filaments are from the same polymerized stock. For different capping protein concentrations, all samples use capping protein from the same aliquot.

### Microscopy

Samples are imaged using a spinning disk confocal (Nikon) equipped with a CMOS camera (Andor) and 60× 1.2NA objective (Nikon). Samples are illuminated using a 491 or 561 nm laser (Cobolt). The polarization images were acquired on a home built LC-Polscope microscope constructed by Rudolph Oldenbourg at Marine Biological Laboratory in Woods Hole, MA [23]

### Image analysis

Droplet aspect ratio is calculated from droplet shape parameters extracted through ImageJ’s built-in Analyze Particles function [24,25]. Images are thresholded and through visual inspection droplets in the process of coalescing, or where thresholding visibly alters the shape, are excluded. To extract the major and minor axes lengths, droplet shape is approximated as an ellipsoid. Droplet shape classification as tactoids or ellipsoids is determined through visual inspection.

To estimate the amount of actin partitioning into droplets, the relative intensities inside to outside the droplet is calculated from a dark subtracted image. Dark is defined as the average intensity without illumination, representing the camera dark levels. The average intensity inside of droplets is compared to the intensity within a ring between 1.4 and 3.2 μm from the droplet border, excluding anywhere within 1.4 μm of another droplet. The ratio between these values is calculated separately for each droplet to account for difference in illumination across the field of view. This ratio is then averaged across all measured droplets.

### Bipolar Tactoids Model

The natural shape for a nematic droplet with strong surface anchoring is a bipolar tactoid [17]. In the bipolar geometry, the tactoid shape is the surface of revolution of an arc of a circle of radius *R* that subtends an angle 2*α* at the center of the circle. *R* and *α* correspond to the size scale and tip angle of the tactoid. The latter specifies the shape in terms of the aspect ratio, *sinα*/(1 − *cosα*). The mechanical free energy of the tactoid can then be written in terms of these parameters as [26]

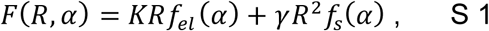

where the nematic elastic energy from the splay and bending of the curved director lines depends on the elastic constant, *K*, and the shape parameter, *f*_*el*_ (*α*) = *π*(−7*α cosα* − 7*sinα* + *α*^2^*sinα*); the surface energy depends on the surface tension, *γ*, and the parameter representing the surface area of the tactoid, *f*_*s*_ (*α*) = 4*π* sin *α*− *αcosα*. Here, we use the equal Frank elastic constant approximation for bend and splay. For a given volume of the droplet, the size and length scale are not independent variables, but related through: 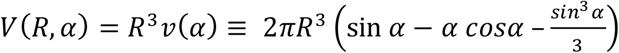. The free energy in Eq. S1 can be re-expressed in terms of the tactoid volume and tip angle, by using 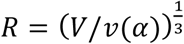, and expressed as a nondimensional free energy,

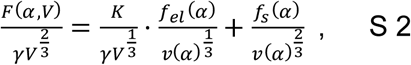

which depends on the tactoid angle and a nondimensional parameter, 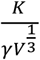, expressing the relative importance of the bulk nematic and surface tension energies. The equilibrium shape, and therefore aspect ratio, of the tactoid of a given volume, *V*, is found by numerically minimizing the nondimensional free energy in Eq. S2 with respect to *α*. We then plot curves for the equilibrium aspect ratio vs. tactoid size (in terms of the cross-sectional area which scales as *V*^2/3^) for different values of the length scale, *K*/*γ*, and compare these to the corresponding experimentally measured aspect ratio vs. area data. These are shown in Fig. 5C. By inspection, we choose three different curves for three different *K*/*γ* values that best describe and bound the data set obtained from averaging over the aspect ratio and area measurements of populations of tactoids. We note that not all tactoids in the same experiment have the same material properties since they have different concentrations of actin with FUS. We thus obtain a range of *K*/*γ* values for each experiment at a different capping protein concentration as shown in Fig. 5D.

**Figure 1.**
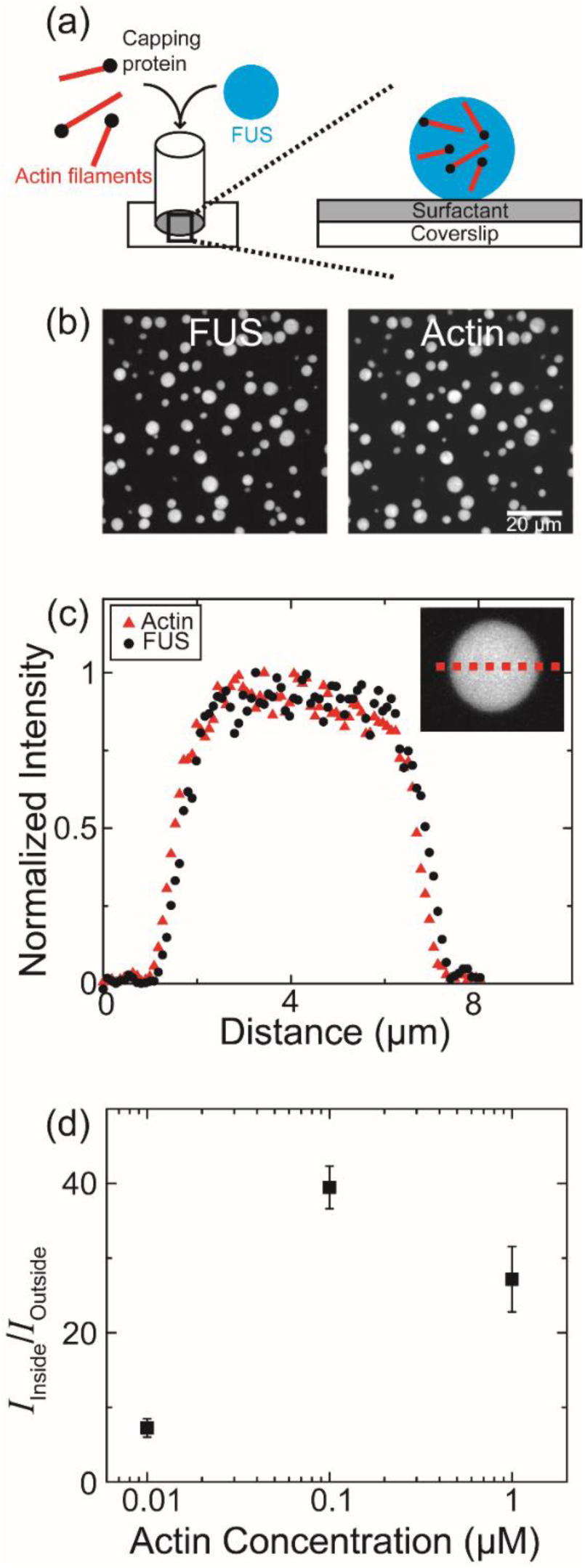
Actin and FUS form composite droplets (a) Schematic of experimental setup. The protein FUS mixes with short, pre-polymerized actin filaments to form composite droplets which sediment to a surfactant passivated layer at the bottom of the sample chamber. (b) Images of composite droplets through fluorescence microscopy of FUS (left) and actin (right). Scale bar is 20 μm (c) Intensity across the midplane of a droplet (*inset*, red dashed line) for FUS (black circles) and actin (red triangles). Intensities are normalized by their maximum values. (d) Intensity of actin in droplets relative to in the solution as a function of actin concentration. Error bars are standard deviation between droplets. In all panels, the data shown are droplet samples composed of 4.4 μM FUS and 1 μM actin with *L* ∼ 90 nm.

**Figure 2.**
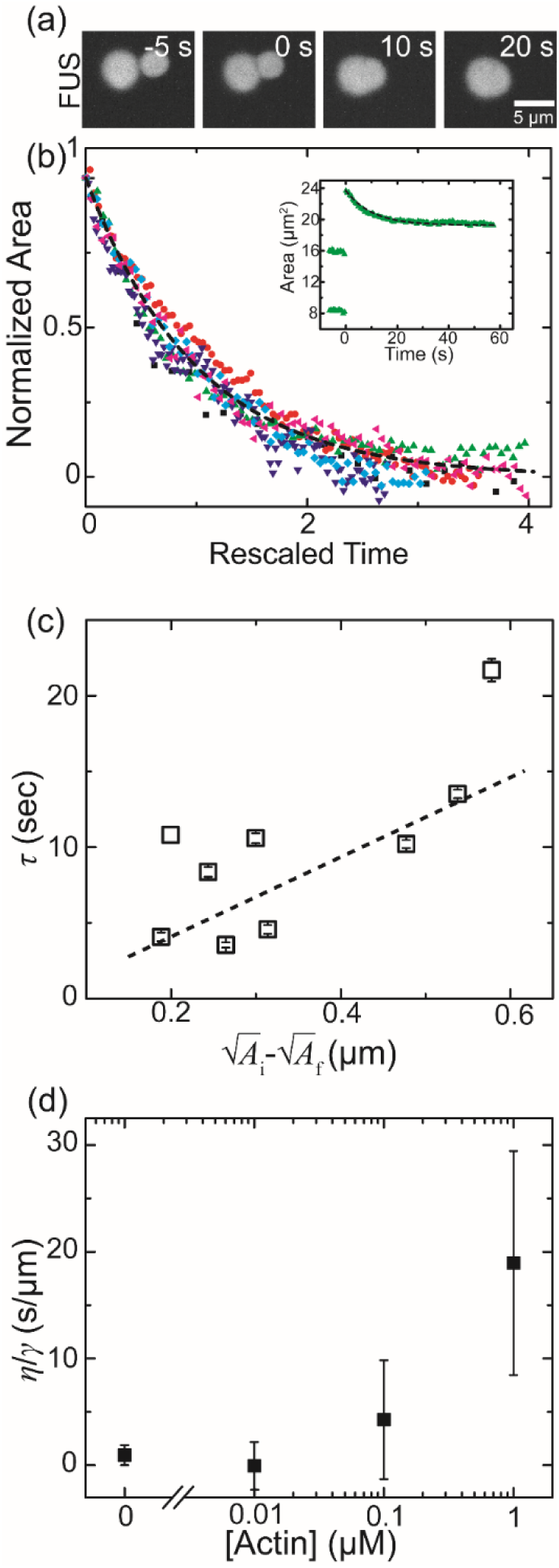
Droplets are liquid with actin dependent properties (a) Fluorescence microscopy images of (FUS-labeled) droplets coalescing over time. Scale bar is 5 μm. (b) *Inset:* Area over time for a single coalescence event (green circles) is fit by a single exponential (black dashed line). Normalized area for 6 different droplet coalescence events with time rescaled by the timescale *τ*. Black dashed line indicates a single exponential, *e*^−*t*^. Data shown is for droplets composed of 4.4 μM FUS and 1 μM actin with *L* ∼ 90 nm. (c) Dependence of the characteristic coalescence time, *τ*, on the coalescence length scale, defined as the difference of the square roots of the final and initial areas. Dashed line indicates a linear fit to the data. Data shown are droplet samples composed of 4.4 μM FUS and 1 μM actin. (d) The ratio of the viscosity to surface tension, given by the slope of the linear fit, such as in part (c), as a function of actin concentration. Error bars are the standard error from the linear fit.

**Figure 3.**
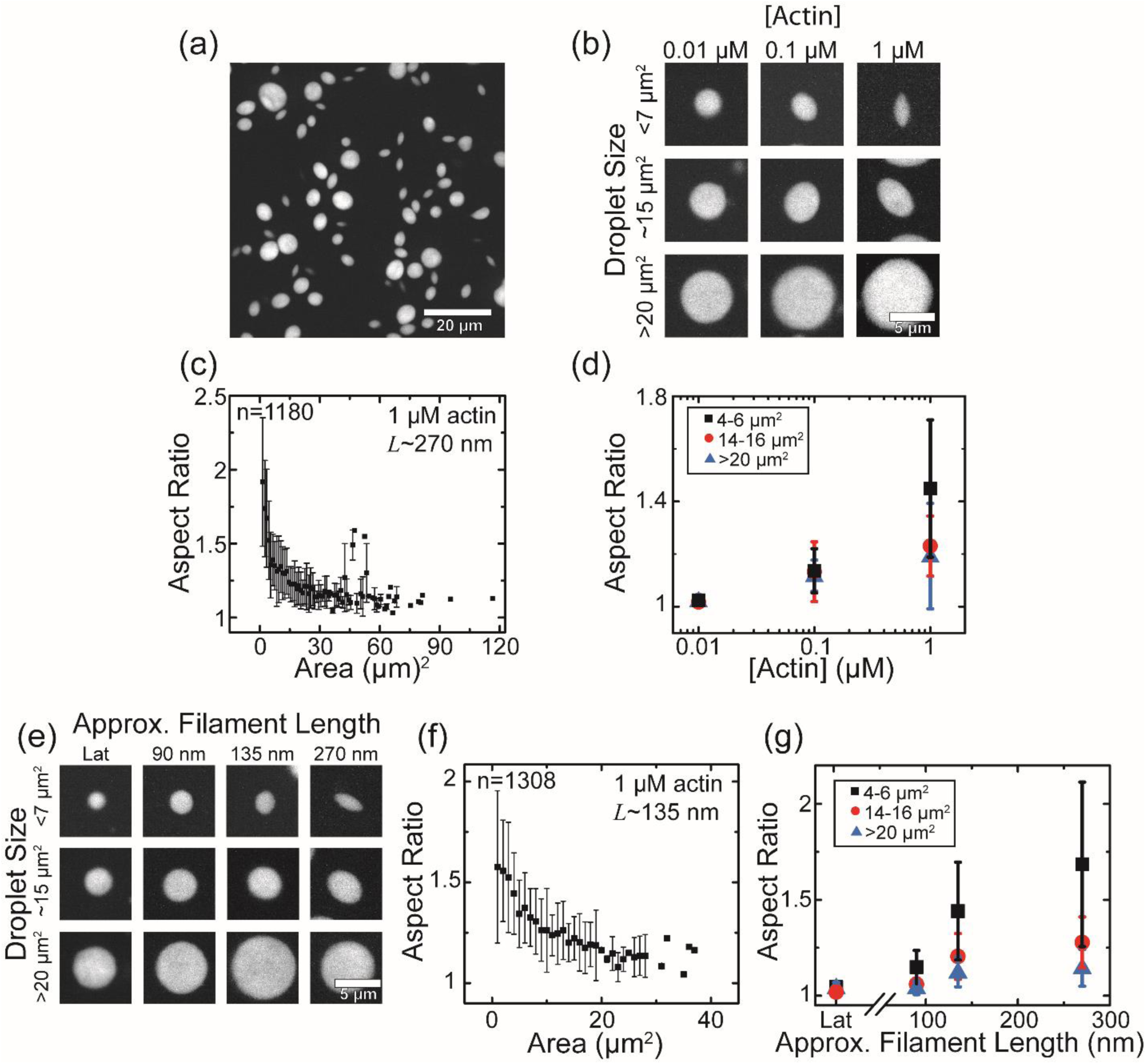
Actin filaments elongate droplets. (a) Fluorescence microscopy image of FUS-labeled droplet samples containing 1 μM actin with *L* ∼ 270 nm. Scale bar is 20 μm. (b-d) Higher actin concentration elongates droplets. All data shown for droplets containing filaments with *L* ∼ 270 nm. (b) Fluorescence images of composite droplets as a function of droplet size and actin concentration. Scale bar is 5 μm. (c) The average aspect ratio of the droplets as a function of cross-sectional area. (d) Actin concentration dependence of average aspect ratio for droplets with areas of 4−6 μm^2^ (black squares), 14−16 μm^2^ (red circles), and > 20 μm^2^ (blue triangles). (e-g) Longer actin filaments elongate droplets. All data shown for droplets with 1 μM actin. (e) Fluorescence microscope images of FUS-labeled droplets as a function of droplet size and actin filament length. In the presence of latrunculin (Lat) which prevents actin polymerization, droplets are spherical regardless of size. Scale bar is 5 μm. (f) Aspect ratio as a function of cross-sectional area for droplets containing actin with *L* ∼ 135 nm. (g) Average aspect ratio as a function of actin filament length. Symbols are the same as in part (d). Error bars represent ± 1 standard deviation between droplets.

**Figure 4.**
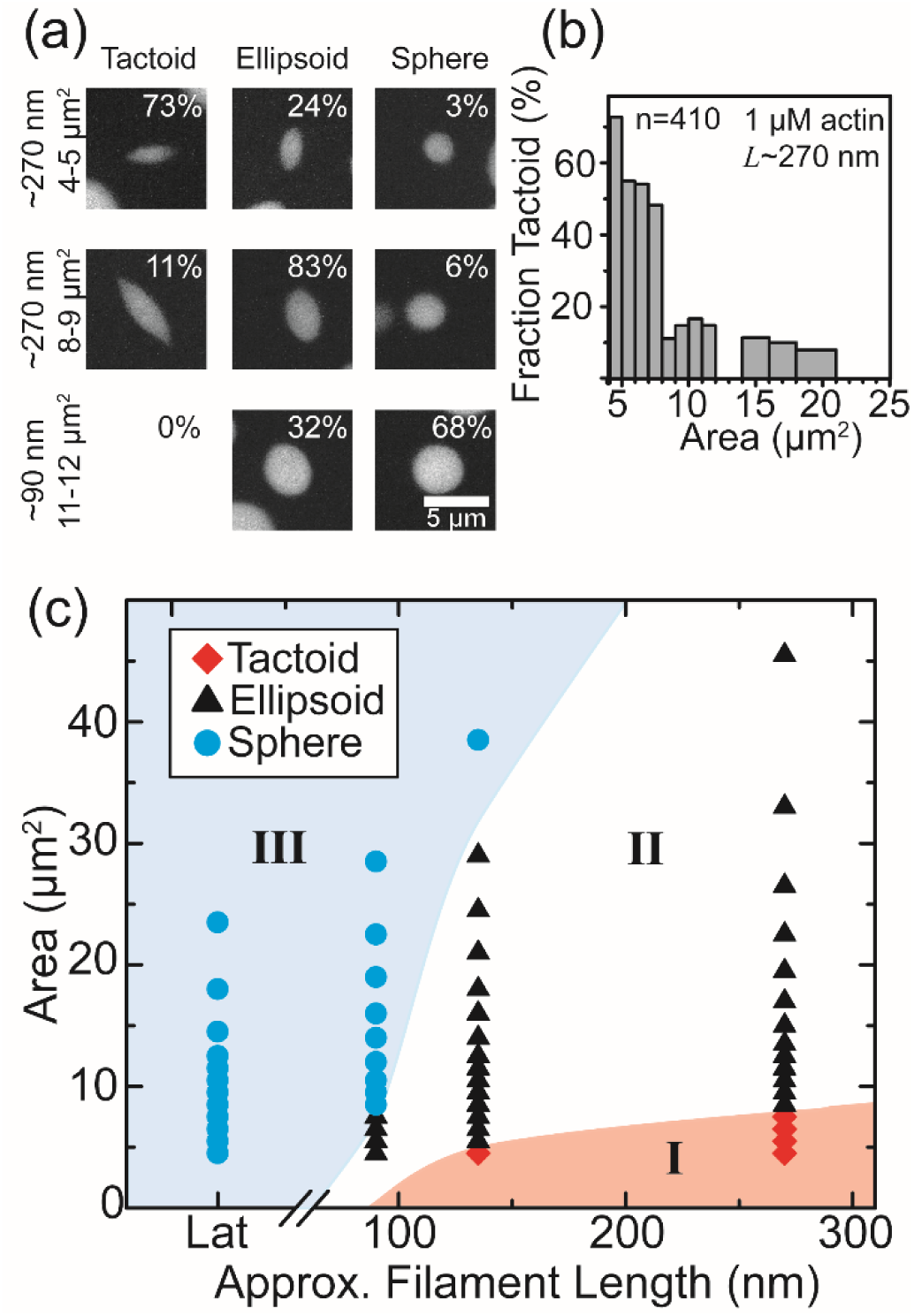
Droplet size and actin length tune the shape of the droplets. (a) Occurrence of droplet shape for three different droplet lengths and corresponding cross-sectional areas. Droplets primarily take on one of three shapes: tactoids with pointed tips (left column), ellipsoids that are elongated but have round tips (center), or spheres (right). (b) Fraction of the droplets that are tactoids as a function of cross-sectional area for droplets containing actin with *L* ∼ 270 nm. Above 25 μm^2^, none of the droplets are tactoids. The shape of droplets under 4 μm^2^ could not be accurately determined. Bar width corresponds to the range of areas included and varies such that each bar represents at least 10 droplets. (c) Phase space of droplet shape as a function of size and filament length, where latrunculin (Lat) indicates unpolymerized actin. We define three regions based on whether droplets are majority (>50%) tactoids (I, red circles), ellipsoids (II, black triangles), or spheres (III, blue diamonds). All samples contain 1 μM actin and 4.4 μM FUS.

**Figure 5.**
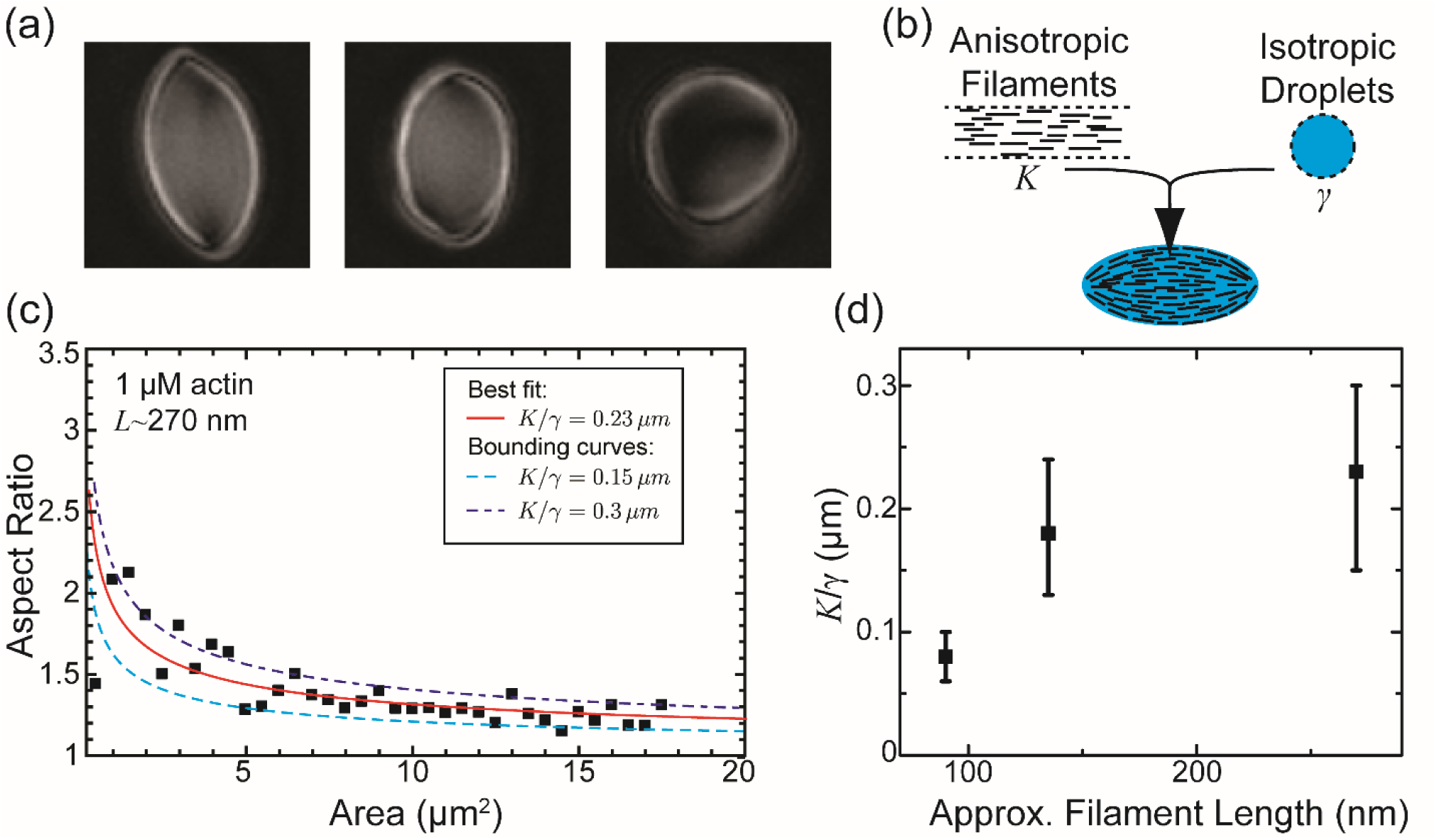
Bipolar model description of droplet shape. (a) LC polscope birefringence images of composite droplets. The dark areas at the poles indicate defects lacking local nematic ordering. (b) Cartoon schematic of droplet shape determinants. Actin filaments contribute a nematic elastic energy, while FUS droplets predominately contribute an isotropic interfacial energy. (c) Theoretical fit (lines) to experimental data (black squares). *K*/γ is extracted from the best fit (red solid line), while the fits that bound the experimental data give the minimum (light blue dashed line) and maximum (dark blue dashed line) values. Data are from samples containing actin with *L* ∼ 270 nm. (d) *K*/γ as a function of actin filament length. Error bars are from the maximum and minimum theoretical fits as shown in part (c). All data are from samples containing 1 μM actin.

## Results and Discussion

To investigate the impact of anisotropic components, such as biopolymer filaments on protein droplets, we sought to form composite droplets of FUS and actin filaments. The RNA-binding protein, FUS, is known to liquid-liquid phase separate into a protein-rich condensed phase upon a reduction in monovalent ion concentration [20]. Here, we form composite FUS-actin droplets by reducing the ambient monovalent salt concentration by an order of magnitude upon dilution, from 500 mM to 45 mM KCl in a solution of FUS-GFP (FUS) and pre-polymerized, fluorescent actin filaments (4.4 μM FUS, 1 μM actin labeled with TMR, 1 mol% capping protein) (Fig. 1a). Using fluorescence microscopy, we observe micrometer-sized condensates enriched with FUS, consistent with previous reports of FUS droplets [20]. Additionally, we find these droplets are also enriched with actin (Fig. 1b). We find that the actin fluorescence uniformly colocalizes with the FUS fluorescence, indicating that these two proteins form composite droplets with apparent homogeneous distribution both actin and FUS (Fig. 1c).

To quantify the partitioning of actin into the droplets, we compare the average actin intensity in the droplet interior, *I*_inside_, to exterior, *I*_outside_. Since fluorescence intensity is proportional to protein concentration, the intensity ratio, *I*_inside_ / *I*_outside_, provides an estimate of the actin concentration inside the droplets relative to in the bulk solution [27,28]. At low actin concentrations (0.01 μM), the intensity of actin inside is about 7 times greater than in the bulk, indicating that the actin filaments preferentially accumulate in the FUS droplets. At higher actin concentrations (0.1 μM −1 μM), the intensity is ∼ 25-40 times greater in the droplets than the bulk solution (Fig. 1d). Thus, actin filaments preferentially incorporate into the FUS droplets across a range of actin concentrations.

Similar to reports of FUS droplets [20], one growth mechanism of composite actin-FUS droplets is coalescence, where two initially separate droplets merge and relax into a new droplet (Fig. 2a and Movie S1) Analyzing the dynamics of coalescence provides an estimate of the relative contributions of two droplet material properties: interfacial tension and viscosity. We measure the droplet cross-sectional area, *A*, as a function of time, *t*, during individual coalescence events (Fig. 2b, *inset*). Two droplets with a total initial cross-sectional area at time of first contact, *A*_*i*_, coalesce into a new droplet which relaxes to a final shape with cross-sectional area, *A*_*f*_. The area decrease is consistent with an exponential decay, which we can extract a characteristic relaxation time, *τ*, from the fit *A*(*t*) = *A*_*f*_ + (*A*_*i*_ − *A*_*f*_)*e*^−*t*/*τ*^. Plotting the normalized area (*A* − *A*_*f*_)/(*A*_*i*_ − *A*_*f*_) against rescaled time *t*/*τ* reveals that coalescence events from various droplet sizes collapse into a single curve that is consistent with an exponential decay (Fig. 2b), indicative of coalescence associated with isotropic fluids [19].

For isotropic fluids, we expect the characteristic relaxation time, *τ*, to scale linearly with the coalescence relaxation length, the difference between the initial and final length, Δ*l*, of the droplet. Here, to account for droplets with shapes that deviate from spherical, we define the relaxation length from the square root of the droplet cross-sectional area such that 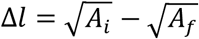. We find that *τ* increases with the difference between the coalescence relaxation length (Fig. 2c). By balancing viscously dissipated mechanical energy against change in the interfacial energy of the coalesced droplet as it relaxes, we see that *τ* depends on the viscosity, *η*, and interfacial tension, *γ*, of the liquid as *τ* ∼ *η*/*γ* · Δ*l* [19]. Thus, the slope of the linear fit in Fig. 2c gives the ratio *η*/*γ*. In Fig. 2d, we find *η*/*γ* increases with actin concentration, suggesting that actin polymer density impacts composite droplet viscosity more than it affects surface tension.

In contrast to FUS droplets which are always spherical [20], composite droplets also adopt a variety of elongated shapes (Fig. 3a). The shape of these droplets is dependent both on the droplet size and actin concentration (Fig. 3b). For low actin concentrations (<0.1 μM), droplets of all assayed cross-sectional areas (between 1 μm^2^ and 32 μm^2^) are spherical. However, for actin concentrations greater than 0.1 μM, we observe non-spherical droplets, particularly in smaller (below ∼15 μm^2^) droplets. To quantify this change in shape, we measure the droplet aspect ratio as a function of cross-sectional area over a range of actin concentrations. For 1 μM actin, elongated shapes are observed for droplet sizes smaller than 20 μm^2^; as the droplet size increases, the average aspect ratio approaches 1 (Fig. 3c). To quantitatively compare the size and actin concentration dependence of droplet shape, we plot average aspect ratio for small (4-6 μm^2^), medium (14-16 μm^2^) and large (>20 μm^2^) droplets. We find the average aspect ratio decreases as droplet size increases, with the trend most pronounced at the highest actin concentration (1 μM). Furthermore, the average aspect ratio increases with actin concentration (Fig. 3d). In contrast to the elongated droplets observed at higher concentrations, for the lowest actin concentration (0.01 μM), droplets of all size have an aspect ratio ∼1. This strongly suggests that the high density of filaments within the drops underlie the observed differences in aspect ratio.

We therefore hypothesize that the actin filaments elongate droplet shape by inducing nematic order. The resulting nematic elasticity competes with the droplet interfacial tension, which constrains pure FUS droplets to be spherical. To test whether the filamentous form of actin, rather than the protein incorporation, is causing the droplet elongation, we form composite droplets in the presence of the drug latrunculin (Lat) which prevents actin polymerization into filaments. In this case, all droplets are spherical (Fig. 3e). Additionally, previous work has shown that nematic elastic energy scales with the aspect ratio of the rod-like constituents [12,13]. We thus hypothesize that filament length impacts droplet shape. To systematically study the impact of filament length on droplet shape, we modify the actin filament length through the amount of capping protein [29]. As the concentration of capping protein is increased from 1 mol% to 3 mol%, the average length of the actin filaments is expected to decrease from ∼270 nm to ∼90 nm. We find that, at 1 μM actin, droplet shape varies with average actin filament length, *L*, with longer filaments leading to more elongated droplets (Fig. 3e). For *L* ∼ 135 nm, smaller (<20 μm^2^) droplets are elongated, with the average aspect ratio decreasing with cross-sectional area (Fig. 3f). In contrast, for the shortest actin filaments tested, *L* ∼ 90 nm, droplets of all sizes have aspect ratios ∼1 (Fig. 3e,g). Notably, for the smallest droplet sizes (4-6 μm^2^), the average aspect ratio increases as the filament length increases.

For a given droplet size and composition, there is a distribution of three characteristic shapes: elongated tactoids with pointed ends, ellipsoids, and spheres. The two elongated shapes, tactoids and ellipsoids, are distinguished by the local shape at the tips of the long axis, where tactoids tips are sharp cusps while ellipsoids tips are smooth (Fig. 4a). To quantify the prevalence of these shapes, we measure the fraction of droplets that are tactoids as a function of cross-sectional area. We visually distinguish between tactoids and ellipsoids or spheres based on the droplet tips, noting that even droplets with ellipsoid shape are technically tactoids if they are nematic liquid crystal droplets. With 1 μM actin, *L* ∼ 270 nm, droplets with cross-sectional areas less than 8 μm^2^ are majority (>50%) tactoids (Fig. 4b). On the other hand, for larger (>8 μm^2^) droplets, the fraction that are tactoids sharply drops to less than 20%. While this transition from tactoid to ellipsoid and sphere with increased size has been theoretically predicted [17,18], to our knowledge it has not been previously experimentally observed. Based on this observation we classify droplets by the shape which is most frequently adopted for each size and filament length. We define any droplet with aspect ratio <1.1 to be a sphere. While in principal this definition is not mutually exclusive with being a tactoid, we do not observe tactoids with aspect ratios this small. Plotting as a function of filament length and droplet size, we find three regions of phase space (Fig. 4c) based on whether the droplets are primarily (>50%) tactoids (I), ellipsoids (II), or spheres (III). The smallest droplets with longest actin filaments primarily form tactoids (Fig. 4c, *Region* I). Larger droplets are primarily elliptical, while the largest are spheres (Fig. 4c, *Regions* II & III). The critical size at which droplet shape transitions to ellipsoids or spheres increases with longer filaments: at *L*∼90 nm, all but the smallest (>8 μm^2^) sizes are primarily spheres and no droplets are majority tactoids down to the smallest droplets measured (4 μm^2^), while at *L* ∼ 270 nm droplets are primarily ellipsoidal from 8 μm^2^ up to the largest size observed (∼45 μm^2^). Droplets containing purely monomeric actin are spherical at all sizes (Fig. 4c, *Region* III, *Lat*). Thus, the shape of composite droplets can be tuned either by changing the concentration or length of actin filaments.

At high densities (1 μM) of actin, elongated spindle shaped droplets are seen to nucleate and become more spheroidal as they grow (Fig. 3a, Movie S2). This decrease in aspect ratio with droplet size is consistent with the bipolar model of tactoids [17,26] where actin filaments align to form a nematic phase with curved director lines that follow the droplet interface. The internal nematic order and its bipolar director orientation for 1 μM actin-FUS droplets is confirmed by observing the droplets under crossed polarizers where intensity corresponds to local nematic order (Fig. 5a). In particular, the reduced intensity seen at the droplet poles suggest the presence of defects (“boojums”) in the nematic order that are characteristic of bipolar tactoids [14].

The shape of a tactoid with characteristic length *R* results from a competition of an interfacial energy that grows as *γR*^2^ with droplet size, and the bulk nematic elastic energy which scales as *K* · *R*, where *K* is the Frank constant (elastic energy scales as droplet volume times the square of the curvature of the director lines, *R*^3^ · *R*^−2^ ∼ *R*) (Fig. 5b). Comparing the two energy scales gives us a characteristic nematic distortion length scale, *K*/*γ*. For increasing droplet size, the interfacial energy becomes progressively more dominant than the bulk elastic energy, resulting in more spherical shapes, that is, lower aspect ratios, that become nearly spherical when *R* ≫ *K*/*γ*. This behavior is seen for 1 μM actin-FUS tactoids (Fig. 4c). We expect, in principle, small enough droplets to be homogeneous [14,16,17], but this transition may occur at droplet sizes below our experimental resolution. Using the bipolar geometry, we numerically minimize the scaled form of the free energy expression to yield expected aspect ratios for given droplet size, measured as average cross-section area in the experiment. In this bipolar model, we expect that the aspect ratio for a given droplet size depends only on one constant: the distortion length scale, *K*/*γ*. Comparing the experimental data to the model then lets us estimate *K*/*γ* for the droplets (Fig. 5c). While for pure actin tactoids, this length scale is estimated to be a few microns [19], here we find that for FUS-actin composite droplets, *K*/*γ* is only hundreds of nanometers. This decrease is consistent with the expectation that presence of FUS increases the interfacial tension contribution but not the nematic elasticity. Interestingly, we find that *K*/*γ* increases with average actin filament length (Fig. 5d), consistent with theoretical predictions and previous experiments in actin nematics that show *K* scaling with filament contour length [12,13]. Intuitively, shorter filaments align more easily along the curved droplet surface. This lower elastic distortion energy cost of filament alignment corresponds to lower *K*.

For low densities (0.01 μM) of actin, the resulting droplets are spherical for all sizes, as is expected of pure FUS droplets. This points to the absence of a nematic phase at such low actin concentration. The Onsager theory does in fact predict a critical density of rods above which they align purely on entropic grounds [9,10]. The critical volume fraction at which nematic order occurs scales inversely as the aspect ratio of the filaments, *L*/*D*, where *D* ∼ 8 nm is the diameter of the constituent actin filaments. This qualitatively explains the observed tendency to form elongated droplets at higher actin concentration as well as at longer average actin filament length. At intermediate densities (0.1 μM), the droplets are slightly elongated and appear ellipsoidal, but their aspect ratio does not depend appreciably on droplet size. They also lack the characteristic pointed ends of a bipolar tactoid. We speculate that this concentration of actin induces some nematic order resulting in a slight anisotropy of the physical properties of the droplet. This results in an ellipsoidal instead of a spherical droplet, while not contributing sufficient nematic order required for the characteristic tactoid shape.

## Conclusions

Here we find that actin filaments spontaneously partition into FUS droplets. Since FUS and actin have no known specific biochemical interaction, this suggests the complexation is driven by non-specific protein-protein interactions such as charge or hydrophobicity. Partitioning of filaments induces anisotropy in otherwise isotropic condensates. While it is well appreciated that modifying droplet macromolecular components and their interactions can influence its mechanical properties from solid-like to liquid-like [7], here we tune its anisotropic properties while maintaining a liquid phase. Moreover, by partitioning actin filaments into a significantly reduced volume, this provides a new route for forming liquid crystal droplets. Future work to understand how the nature of macromolecular component interactions control their miscibility, spatial organization and mechanics will be an exciting area of inquiry.

One consequence of liquid crystallinity is that competing effects of elasticity with interfacial tension give rise to diverse droplet shapes and internal structure. For example, we have shown experimentally the transition between spherical and tactoid-shaped droplets which has been theoretically predicted [17]. This shape change inherently results in varying the surface area to volume ratio, which could be a mechanism to dynamically tune partitioning or other interface-mediated activity. Furthermore, these shapes reflect changes in the spatial organization of the filaments across the droplet [18]. This internal structure can be used as a template for droplet-scale spatial structure [30–32]. Thus, composite condensates offer a promising means to understand and design reconfigurable materials where the interfacial and elastic phases are orthogonally tuned.

Phase separation is well appreciated as a mechanism of intracellular organization [33]. We speculate that the myriad of biopolymers found within the cytoplasm could spontaneously partition to these condensates to form similar composite drops in vivo. One outstanding example speculated to form a liquid crystalline phase is that of the mitotic spindle [34]. Recently, evidence for a “spindle matrix” comprised of protein-rich condensate around microtubule filaments [35] suggests an analog of the composite we observe. Finally, the extent to which this may influence intermediate filament and F-actin organization is unknown, but has potential implications for neurodegenerative diseases [36], and control of cytoskeletal signaling [37].

## Acknowledgements

We thank Anthony Hyman for insightful conversations at the HHMI HCIA Summer Institute and for generous gifts of reagents. We thank Rudolph Oldenbourg for generous assistance in obtaining the polarization images. We appreciate the many discussions and interactions throughout the collaborative community at the HHMI HCIA Summer Institute and the Marine Biological Laboratory that inspired this research. This work was supported by the 2017 HHMI HCIA Summer Institute and partially supported by the University of Chicago MRSEC, supported by NSF 1520709. MLG also acknowledges support from NSF DMR-1905675 and ARO MURI W911NF1410403.

---

**Movie S1.** Coalescence of two droplets composed of 1 μM actin with *L* ∼ 90 nm and 4.4 μM FUS. The time when two droplets initially make contact at the beginning of coalescence is defined as *t* = 0 s. Time is in min:sec.

**Movie S2.** Droplet nucleation and growth. Droplets are composed of 1 μM actin with *L* ∼ 135 nm and 4.4 μM FUS. The time at which FUS and actin are initially mixed together is defined as *t* = 0 s. Time is in min:sec.

## References

[1] A. B. Marciel, E. J. Chung, B. K. Brettmann and L. Leon, Adv. Colloid Interface Sci., 2017, 239, 187–198.

[2] C. P. Brangwynne, P. Tompa and R. V. Pappu, Nat. Phys., 2015, 11, 899–904.

[3] T. K. Lytle, L. W. Chang, N. Markiewicz, S. L. Perry and C. E. Sing, ACS Cent. Sci., 2019, 5, 709–718.

[4] S. F. Banani, A. M. Rice, W. B. Peeples, Y. Lin, S. Jain, R. Parker and M. K. Rosen, Cell, 2016, 166, 651–663.

[5] L. Li, S. Srivastava, M. Andreev, A. B. Marciel, J. J. de Pablo and M. V. Tirrell, Macromolecules, 2018, 51, 2988–2995.

[6] X. Wang, J. Lee, Y. W. Wang and Q. Huang, Biomacromolecules, 2007, 8, 992–997.

[7] D. V. Krogstad, N. A. Lynd, S. H. Choi, J. M. Spruell, C. J. Hawker, E. J. Kramer and M. V. Tirrell, Macromolecules, 2013, 46, 1512–1518.

[8] M. Antonov, M. Mazzawi and P. L. Dubin, Biomacromolecules, 2010, 11, 51–59.

[9] L. Onsager, Ann. N. Y. Acad. Sci., 1949, 51, 627–659.

[10] P. G. de Gennes and J. Prost, The physics of liquid crystals, Oxford University Press, Oxford, 1995.

[11] T. Odijk, Liq. Cryst., 1986, 1, 553–559.

[12] J. P. Straley, Phys. Rev. A, 1973, 8, 2181–2183.

[13] R. Zhang, N. Kumar, J. L. Ross, M. L. Gardel and J. J. de Pablo, Proc. Natl. Acad. Sci. U. S. A., 2018, 115, E124–E133.

[14] V. Jamali, N. Behabtu, B. Senyuk, J. A. Lee, I. I. Smalyukh, P. van der Schoot and M. Pasquali, Phys. Rev. E, 2015, 91, 042507.

[15] P. W. Oakes, J. Viamontes and J. X. Tang, Phys. Rev. E, 2007, 75, 061902.

[16] M. Bagnani, G. Nyström, C. De Michele and R. Mezzenga, ACS Nano, 2019, 13, 591–600.

[17] P. Prinsen and P. van der Schoot, Phys. Rev. E, 2003, 68, 11.

[18] R. M. W. Van Bijnen, R. H. J. Otten and P. Van Der Schoot, Phys. Rev. E, 2012, 86, 1–12.

[19] K. L. Weirich, S. Banerjee, K. Dasbiswas, T. A. Witten, S. Vaikuntanathan and M. L. Gardel, Proc. Natl. Acad. Sci., 2017, 114, 2131–2136.

[20] A. Patel, H. O. Lee, L. Jawerth, S. Maharana, M. Jahnel, M. Y. Hein, S. Stoynov, J. Mahamid, S. Saha, T. M. Franzmann, A. Pozniakovski, I. Poser, N. Maghelli, L. A. Royer, M. Weigert, E. W. Myers, S. Grill, D. Drechsel, A. A. Hyman and S. Alberti, Cell, 2015, 162, 1066–1077.

[21] J. A. Spudich and S. Watt, J. Biol. Chem., 1971, 246, 4866–4871.

[22] S. Palmgren, P. J. Ojala, M. A. Wear, J. A. Cooper and P. Lappalainen, J. Cell Biol., 2001, 155, 251–60.

[23] R. Oldenbourg, Nature, 1996, 381, 811–812.

[24] C. A. Schneider, W. S. Rasband and K. W. Eliceiri, Nat. Methods, 2012, 9, 671–675.

[25] J. Schindelin, I. Arganda-Carreras, E. Frise, V. Kaynig, M. Longair, T. Pietzsch, S. Preibisch, C. Rueden, S. Saalfeld, B. Schmid, J.-Y. Tinevez, D. J. White, V. Hartenstein, K. Eliceiri, P. Tomancak and A. Cardona, Nat. Methods, 2012, 9, 676–682.

[26] A. V. Kaznacheev, M. M. Bogdanov and S. A. Taraskin, J. Exp. Theor. Phys., 2002, 95, 57–63.

[27] S. F. Banani, A. M. Rice, W. B. Peeples, Y. Lin, S. Jain, R. Parker and M. K. Rosen, Cell, 2016, 166, 651–663.

[28] P. M. McCall, S. Srivastava, S. L. Perry, D. R. Kovar, M. L. Gardel and M. V. Tirrell, Biophys. J., 2018, 114, 1636–1645.

[29] A. Weeds and S. Maciver, Curr. Opin. Cell Biol., 1993, 5, 63–69.

[30] K. L. Weirich, K. Dasbiswas, T. A. Witten, S. Vaikuntanathan and M. L. Gardel, Proc. Natl. Acad. Sci., 2019, 116, 201814854.

[31] P. Poulin, H. Stark, T. C. Lubensky and D. A. Weitz, Science, 1997, 275, 1770–1773.

[32] X. Wang, D. S. Miller, J. J. De Pablo and N. L. Abbott, Soft Matter, 2014, 10, 8821–8828.

[33] A. A. Hyman, C. A. Weber and F. Jülicher, Annu. Rev. Cell Dev. Biol., 2014, 30, 39–58.

[34] J. Brugués and D. Needleman, Proc. Natl. Acad. Sci., 2014, 111, 18496–18500.

[35] H. Jiang, S. Wang, Y. Huang, X. He, H. Cui, X. Zhu and Y. Zheng, Cell, 2015, 163, 108–122.

[36] N. J. Cairns, V. M.-Y. Lee and J. Q. Trojanowski, J. Pathol., 2004, 204, 438–449.

[37] X. Su, J. A. Ditlev, E. Hui, W. Xing, S. Banjade, J. Okrut, D. S. King, J. Taunton, M. K. Rosen and R. D. Vale, Science, 2016, 352, 595–599.

